# Coupling between sterol and sphingolipid structure in ordered membrane domains

**DOI:** 10.64898/2026.04.01.715929

**Authors:** Israel Juarez-Contreras, Hyesoo Kim, Itay Budin

## Abstract

A hallmark of eukaryotic membranes is the pairing of lineage-specific sterols with characteristic sphingolipid species. Mammalian cell membranes are enriched in both cholesterol and long-chain sphingolipids like sphingomyelin, whereas fungi synthesize ergosterol and very long-chain sphingolipids with sugar-containing head groups. It has been proposed that these two lipid classes co-evolved to support membrane structure and organization. Here we investigated how sterol structure and sphingolipid chain length together control membrane order and phase behavior. In the yeast *Saccharomyces cerevisiae*, loss of very long-chain C26 sphingolipids disrupted formation of liquid-ordered (*L*_o_) domains in the vacuole membrane. Similarly, substitution of ergosterol synthesis for that of cholesterol also prevented vacuole *L*_o_ domains. To determine a possible physical basis of these effects, we investigated synthetic membranes of defined composition containing either ergosterol or cholesterol and sphingomyelin with different chain lengths. In membranes containing egg sphingomyelin with C16 chains, ergosterol only sparsely supported *L*_o_ domains, in contrast to cholesterol. Membranes containing sphingomyelin with C26 chains displayed a different pattern. Cholesterol mixtures were largely homogeneous across most compositions, with only a limited region that supported fluid domains. Ergosterol mixtures exhibited a distinct compositional window that supported fluid domains positioned between regimes of uniform membranes and gel phases. This window corresponded to stoichiometric changes in the vacuole as it phase-separates during nutritional restriction. Measurements of membrane order showed that cholesterol strongly increased membrane packing compared to ergosterol in membranes containing egg sphingomyelin, whereas this difference was lost in membranes containing C26 sphingomyelin. The results suggest that sphingolipid chain length can tune sterol interactions needed for membrane organization.

**Significance:** Membrane phase separation into coexisting ordered and disordered fluid domains has largely been investigated using characteristic mammalian lipid components, cholesterol and long-chain saturated lipids like sphingomyelin. Under nutrient limitation, vacuole membranes in yeast organize into micron-scale domains that are important for their physiology. Compared to mammals, yeast synthesize an alternative sterol, ergosterol, and sphingolipids with very long-chains. We show that vacuole membrane domains are sensitive to both these features, which also show preferential interactions in liposomes that support membrane ordering and phase properties. In lipid mixtures containing very long-chain sphingomyelins, stoichiometric regimes that support phase separation of fluid domains are similar to those of the vacuole lipidome under nutrient limitation. This finding supports a model in which sterols and sphingolipids co-evolved to support membrane structure.

## Introduction

Phase separation of membranes provides a potential physical mechanism for lateral organization in living cells and model membrane systems (1, 2). In mixtures of biologically-relevant lipids, membranes can segregate into coexisting liquid phases with distinct physical properties, commonly described as liquid ordered (*L*_o_) and liquid disordered (*L*_d_) phases (3–5). In contrast to solid or gel-like domains, *L*_o_ domains are fluid, and thus *L*_o_/*L*_d_ phase separation achieves lateral organization while maintaining dynamics thought to be required for biological function. Across a range of model membranes, systems that demix into micron-scale fluid phases share a common chemical requirement: the presence of lipids with high-melting temperatures due to their saturated chains, lipids with low melting temperatures due to their unsaturated or branched chains, and sterols like cholesterol (6). Sterols interact with lipid chains, increasing their order, while also preventing the formation of crystalline-like packing of saturated chains that drives solid membrane phases.

While initial studies on *L*_o_/*L*_d_ domains focused on systems containing saturated phospholipids, sphingolipids are the primary saturated lipid species in eukaryotic membranes and can also support *L*_o_/*L*_d_ domains when mixed with sterols and unsaturated phospholipids. Sphingolipids feature largely saturated sphingoid bases and *N*-acyl chains that promote tight lipid packing (7–9) and can specifically interact with sterols through hydrogen bonding (10). Similarly, the ability of sterols to promote ordered membrane phases depends on their chemical structure. Different sterols can induce distinct phase behaviors in model and cellular membranes (11–14). These differences are biologically relevant because sterol and sphingolipid structure vary widely across organisms (1). Mammalian membranes generally contain cholesterol together with sphingomyelin and glycosphingolipids. Of these, sphingomyelin is the most abundant and generally features long *N*-acyl chains (C16-C18). In contrast, fungal membranes largely contain ergosterol and a set of glycosphingolipids that carry very long chains (>C22) (15).

Longer-chain sphingolipids increase bilayer thickness and can introduce hydrophobic mismatch between neighboring lipids, altering lipid packing (16). Very long chains may also extend across the bilayer midplane and promote interdigitation between opposing leaflets (17). Because sterols insert between lipid acyl chains and interact preferentially with saturated lipids, changes in sphingolipid chain length can also alter sterol–lipid packing interactions within the bilayer (18, 19). Studies of cholesterol-sphingomyelin mixtures have shown that sterol interactions with sphingolipid depend on bilayer thickness and acyl-chain packing (20). These previous reports all suggest that variation in sphingolipid chain length may influence membrane phase behavior of sterol-containing membranes.

In yeast, the lipidome of the vacuole membrane remodels during the transition from logarithmic growth to stationary phase, increasing the abundance of both sterols and sphingolipids (21). These changes to the vacuole lipidome correspond to the appearance of membrane domains (22, 23) that are utilized for microautophagy of lipid droplets under nutritional stress (24, 25). Vacuole domains recruit specific proteins, are large, observable by light microscopy, and show the characteristics of *L*_o_ domains (26, 27). Domains are acutely-sensitive to sterol structure (13), which has been proposed to reflect evolutionary optimization of membrane biophysical properties (28). Yeast cells that synthesize either higher- or lower-ordering ergosterol intermediates do not show vacuole domains, corresponding to the capacity of ergosterol’s moderate ordering to support fluid domains in liposomes (13). Vacuole phase separation is also reliant on sphingolipid metabolism and transport, with loss of either preventing domains from forming (21, 29). However, yeast sphingolipids, like inositolphosphoceramide (IPC), are challenging to synthesize or extract from cells and, with few exceptions (30), have not been analyzed in model membranes. As a result, how these sterol–sphingolipid combinations influence membrane phase behavior remains underexplored.

In this study, we examined how sphingolipid acyl chain length influences membrane phase behavior. To isolate the effects of chain length, we generated model membranes containing ergosterol and sphingomyelin species with either relatively short or very long chains, with the latter mimicking those found in fungal membranes. Using giant and large unilamellar vesicles, we compared how these lipids influence liquid–liquid phase separation and membrane packing. We report how sphingolipid chain length modulates sterol-dependent membrane phases and ordering, showing that systems with ergosterol and C26 sphingolipids better mimic the phase behavior of the vacuole membrane as it phase-separates in cells.

## Results

### Formation of vacuole domains depend on both sterol structure and sphingolipid chain length

The yeast vacuole undergoes pronounced membrane remodeling as cells enter stationary phase or are subjected to nutrient stress, producing micron-scale domains that segregate membrane proteins and lipids (21, 23, 31). To examine how sterol structure and sphingolipid chain length contribute to this process, cells were grown to stationary phase under glucose depletion to induce vacuole membrane domain formation and vacuoles were visualized using Pho8-GFP as a liquid-disordered (*L*_d_) marker. Wild-type (WT) vacuoles displayed micron-scale domains from which Pho8-GFP was excluded, consistent with previous observations of phase-separated vacuole membranes (Fig 1C). To test whether yeast’s very long sphingolipid chains specifically contribute to domain formation, we examined mutants defective in sphingolipid elongation. *ELO2* and *ELO3* encode fatty acid elongases required for synthesis of long acyl chains characteristic of yeast sphingolipids. Previous analyses showed that *elo2*Δ mutants produced shortened sphingolipids, but still contained C26 species, whereas e*lo3*Δ mutants completely lack C26 sphingolipids (15). Vacuoles from *elo2Δ* cells retained visible membrane domains, but at reduced frequency compared to wild type (Fig. 1D, Fig. S1A). In contrast, *elo3Δ* vacuoles lacked any detectable domains under these conditions (Fig. 1D, Fig. S1A). Loss of domains in elongation mutants was also accompanied by increased vacuole fragmentation (Fig. S1B). Fragmented vacuoles do not phase-separate, but even the minority of fused vacuoles in e*lo3Δ* cells did not show any detectable domains. These observations indicate that very long chain sphingolipids contribute to the formation of vacuole domains.

**Figure 1.**
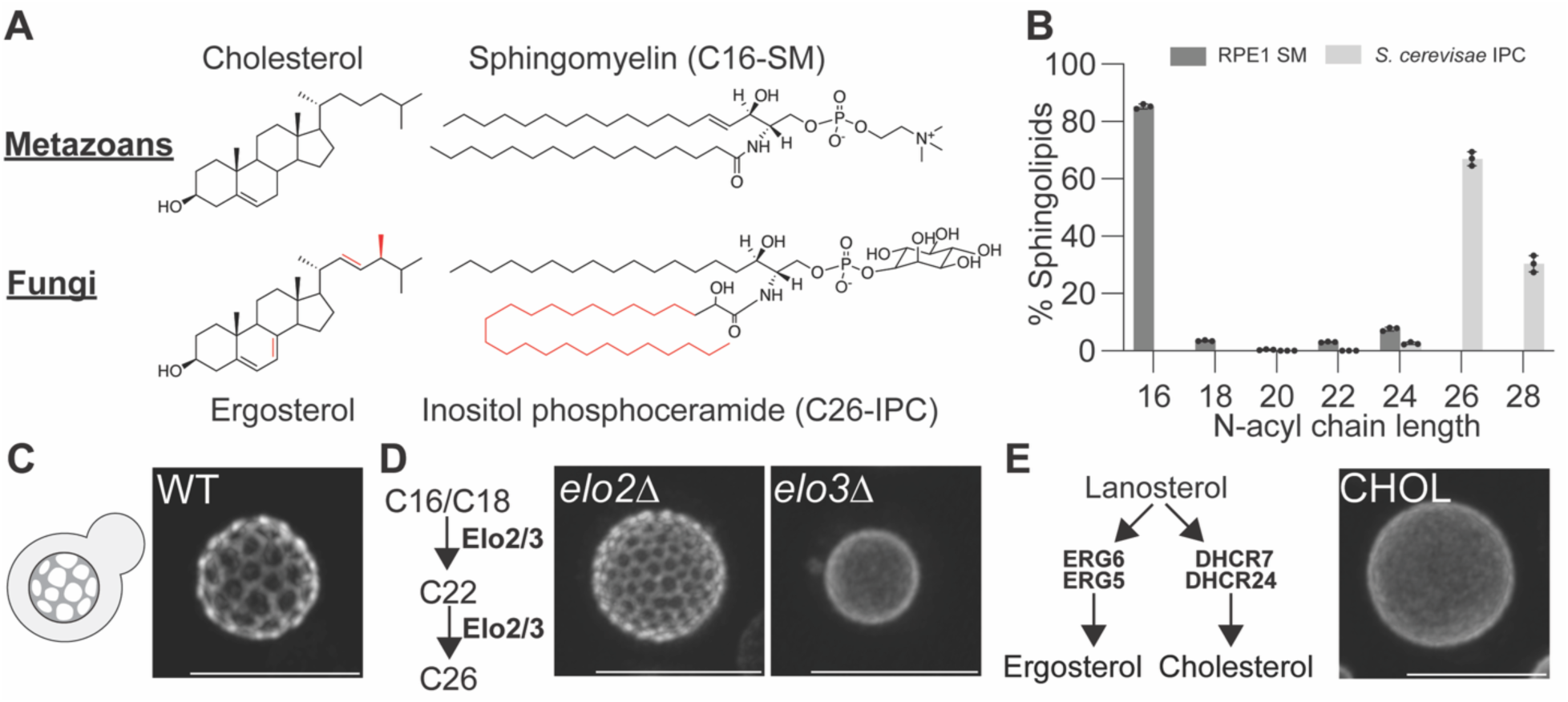
Vacuole membrane domains require ergosterol and very long-chain sphingolipids. (**A**) Canonical sterol-sphingolipid pairs in metazoans and fungi. Metazoan membranes are enriched in cholesterol and C16 sphingomyelin (SM), whereas fungal membranes contain ergosterol and very long chain (C26) inositol phosphoceramide (IPC). Structural features unique to ergosterol and IPC are highlighted in red. (**B**) Sphingolipid (SL) chain length distributions. Retinal pigment epithelial (RPE1) cells are enriched in C16 sphingomyelin, with minimal contribution from longer chains, whereas *Saccharomyces cerevisiae* sphingolipids are dominated by very long chain species (C26-C28). Lipidomic data is replotted from previous studies (21, 36). (**C**) Schematic and representative fluorescence micrograph of vacuole membrane domains in WT yeast, showing micron-scale phase separation. (**D**) Disruption of sphingolipid elongation alters vacuole membrane organization. Sphingolipid acyl chains are extended from C16/C18 to C26 via Elo2 and Elo3. Deletion of *ELO2* or *ELO3* shortens sphingolipid chains and reduces micron-scale vacuole domains, as shown in representative micrographs and quantification in Fig. S1A. In contrast to *elo2Δ* cells, *elo3Δ* cells lack any C26 sphingolipids and show no domains. (**E**) Sterol structure also determines vacuole membrane organization. Lanosterol is a shared precursor in both metazoan and fungal sterol pathways. Ergosterol is synthesized from lanosterol through the ERG pathway, with Erg6 and Erg6 catalyzing key steps that generate fungal-specific structural features. Replacement of these enzymes with the metazoan counterparts DHCR24 and DHCR7 redirects the pathway to produce cholesterol, while upstream steps remain functionally compatible. Cells producing cholesterol instead of ergosterol fail to form vacuole membrane domains, demonstrating a requirement for sterol structural specificity. For C-E, scale bars, 5 µm.

The structure of ergosterol also has a strong influence on the formation of vacuole domains (13). Cholesterol promotes membrane ordering more robustly than ergosterol and also supports *L*_o_/*L*_d_ phase separation. To test if this dynamic applies in the vacuole, we engineered yeast strains to synthesize cholesterol in place of ergosterol. As previously shown (32), substitution of two ergosterol biosynthetic genes (*ERG6* and *ERG5*) with those in cholesterol biosynthesis (DHCR7 and DHCR24) results in formation of cholesterol in yeast (Fig. S2A, S2D) in place of ergosterol (Fig. S2B, S2D). The resulting strain (CHOL) synthesizes cholesterol to the same level as WT cells synthesize ergosterol (Fig. S2C). Despite cholesterol’s propensity to support *L*_o_/*L*_d_ domains in model membranes, vacuoles in CHOL cells showed a completely uniform Pho8-GFP distribution (Fig. 2B). These observations led us to ask if a structural compatibility between sterols and very long chain sphingolipids contributes to vacuole membrane organization. Because loss of Elo3 and ergosterol synthesis induced a synthetic lethality, as previously observed (33), we turned to model membranes to further investigate this question.

**Figure 2.**
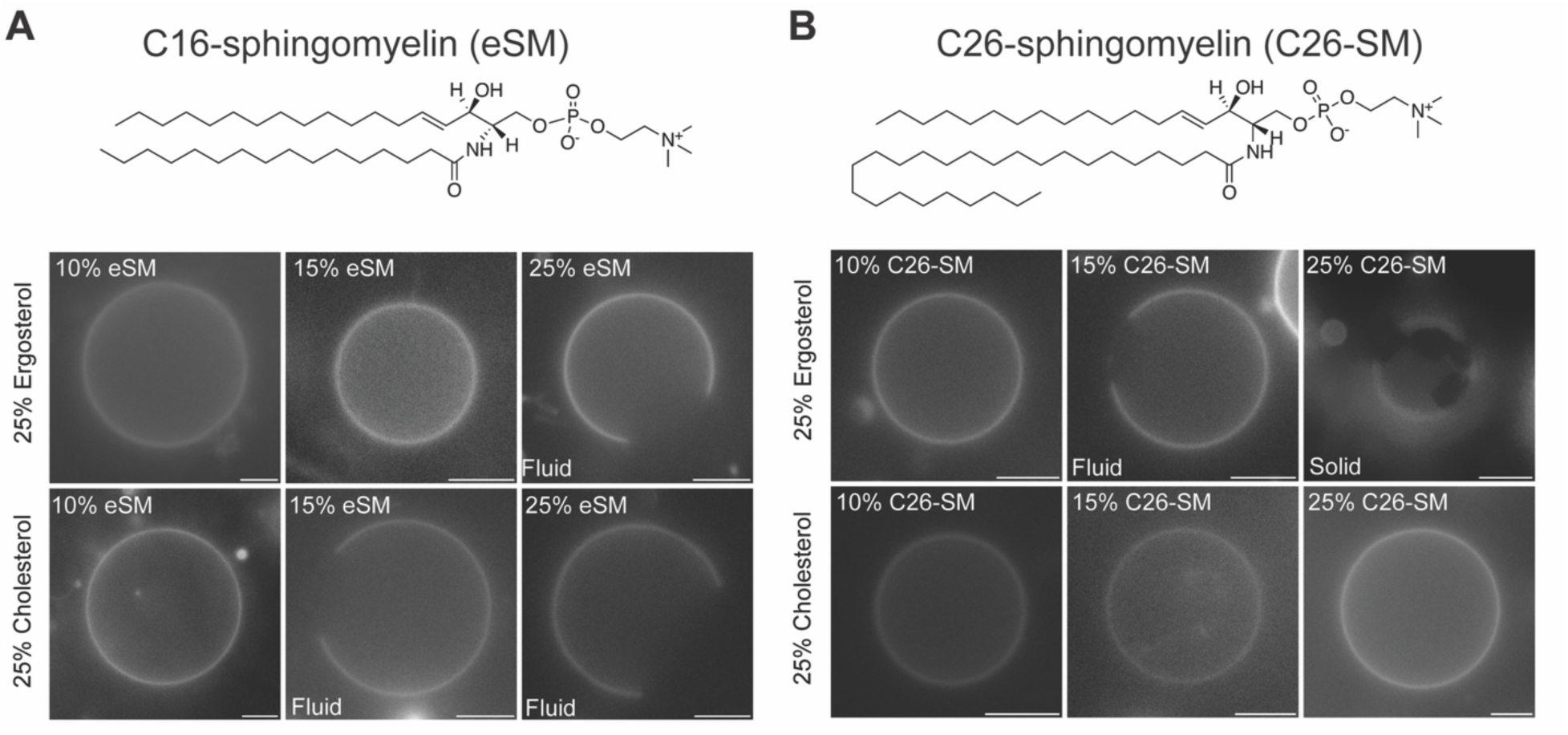
Sphingomyelin chain length tunes membrane phase behavior. (**A**) Representative micrographs of GUVs containing 25% sterol and increasing egg sphingomyelin (eSM). In ergosterol-containing membranes (top), fluid domains are maintained only at 25% eSM, whereas cholesterol-containing membranes (bottom) support fluid domains across a broader eSM range. (**B**) Substitution with very-long-chain C26-SM induces phase separation and solid-like behavior in ergosterol, but not cholesterol. Scale bars, 5 µm.

### Ergosterol and cholesterol induce distinct phase behaviors with C26 sphingomyelin

To test whether sterol structure and sphingolipid chain length directly influence membrane phase behavior, we examined sterol-sphingomyelin mixtures in defined lipid compositions. Previous work has shown that substitution of ergosterol for cholesterol (12, 14) can influence membrane miscibility and ordered domain formation, but variations to sphingolipid structure have been less well explored. Notably, substitution of sphingomyelin in glucosylceramide can cause formation of gel-like domains in place of fluid *L*_o_ ones (34). In yeast sphingolipids differ from those found in mammals with respect to headgroup structure, hydroxylation, and chain length (Fig. 1A, 1B). To isolate the role of the latter, we utilized a very long-chain C26 sphingomyelin (C26-SM), comparing its effects to that of well-studied egg sphingomyelin (eSM) with C16 *N*-acyl chains. C26-SM is not abundant in mammalian lipidomes, which feature C16 and other long-chain sphingomyelins, as well as somewhat longer-chained hexosylceramides (35, 36). It is also not found in yeast that do not synthesize SM. Instead, it serves as a useful synthetic model for investigating the C26 *N*-acyl chains that dominate yeast sphingolipids.

To explore sterol-sphingolipid interactions, we prepared giant unilamellar vesicles (GUVs) containing 25 mol% of either ergosterol or cholesterol and prepared with increasing fractions (10-25%) of either eSM or C26-SM. This concentration range was designed to sample compositions commonly used to investigate *L*_o_/*L*_d_ phases, as well as modeling the sphingolipid changes to yeast vacuole membranes from ∼15 mol % total sphingolipids (21, 37). The remaining lipids were the unsaturated phospholipid di-oleoyl phosphatidylcholine (DOPC) as a low melting temperature component and trace quantities of Texas Red-labeled dihexadecanoyl phosphatidylethanolamine (DHPE) as a *L*_d_ marker (Fig. 2). Across these compositions, three membrane behaviors were observed: homogeneous fluid membranes, liquid-liquid coexistence (*L*_o_/*L*_d_) of ordered domains that are depleted in Texas Red-DHPE, and solid-like domains that are depleted in Texas Red-DHPE and do not fuse with one another. Similar regimes have been described in ternary sterol-sphingolipid-phospholipid membranes, where differences in lipid packing produce fluid-fluid coexistence or transitions into gel-like states depending on composition (3, 38).

Membranes containing eSM displayed clear differences between the two sterols. Ergosterol-containing membranes remained largely homogeneous at 10% and 15% eSM, with liquid-liquid coexistence appearing only at 25% eSM (Fig. 2A). In contrast, cholesterol-containing membranes exhibited liquid-liquid coexistence at both 15% and 25% eSM. Cholesterol is well known to promote *L*_o_/*L*_d_ coexistence in sphingomyelin-containing membranes (4, 38), and comparative phase diagrams of cholesterol-sphingomyelin mixtures show broad regions of fluid phase separation across similar lipid compositions (39). These observations suggest that cholesterol more readily promotes ordered lipid assemblies when sphingolipid chains are relatively short. This effect could be explained by cholesterol’s capacity to condense acyl chains to a greater extent than ergosterol (8, 40, 41).

Membranes containing C26-SM displayed a different pattern than those with eSM (Fig. 2B). The cholesterol systems were homogeneous across the three C26-SM compositions, indicating that the longer-chain sphingomyelin inhibits phase separation in cholesterol-containing membranes. In contrast, the ergosterol system with 15 mol% C26-SM, mimicking vacuole membrane levels, showed liquid-liquid coexistence. Reduced C26-SM (10 mol%) levels led to the disappearance of domains, while increased (25 mol%) levels caused the formation of solid domains. Increasing sphingomyelin acyl chain length therefore alters how each sterol influences membrane organization.

### Systems very long-chain sphingomyelin and ergosterol mimic the induction of vacuole domain

The GUV observations suggested that sphingomyelin chain length alters membrane phase behavior. To examine this effect systematically, we constructed ternary phase diagrams from a range of experimentally sampled GUV compositions (Fig. 3), again classifying each composition as homogeneous, showing liquid-liquid domain coexistence, or exhibiting solid-like domains based on the segregation of Texas Red-DHPE and their capacity to coalesce into larger domains. For each system, at least 30 GUVs were assessed and phase behavior was assigned to the most abundant features observed (Fig. S3-S5). Similar previous studies with ergosterol and saturated PC lipids (42, 43), membranes containing ergosterol, eSM, and DOPC displayed a limited region of liquid-liquid coexistence (Fig. 3A, Fig. S3). This coexistence regime was in stark contrast to the large area of *L*_o_/*L*_d_ coexistence widely reported in phase diagrams of cholesterol/eSM/DOPC (39, 44). Notably, several compositions that fall squarely within the liquid-liquid coexistence region of cholesterol systems instead produced solid domains with ergosterol.

**Figure 3.**
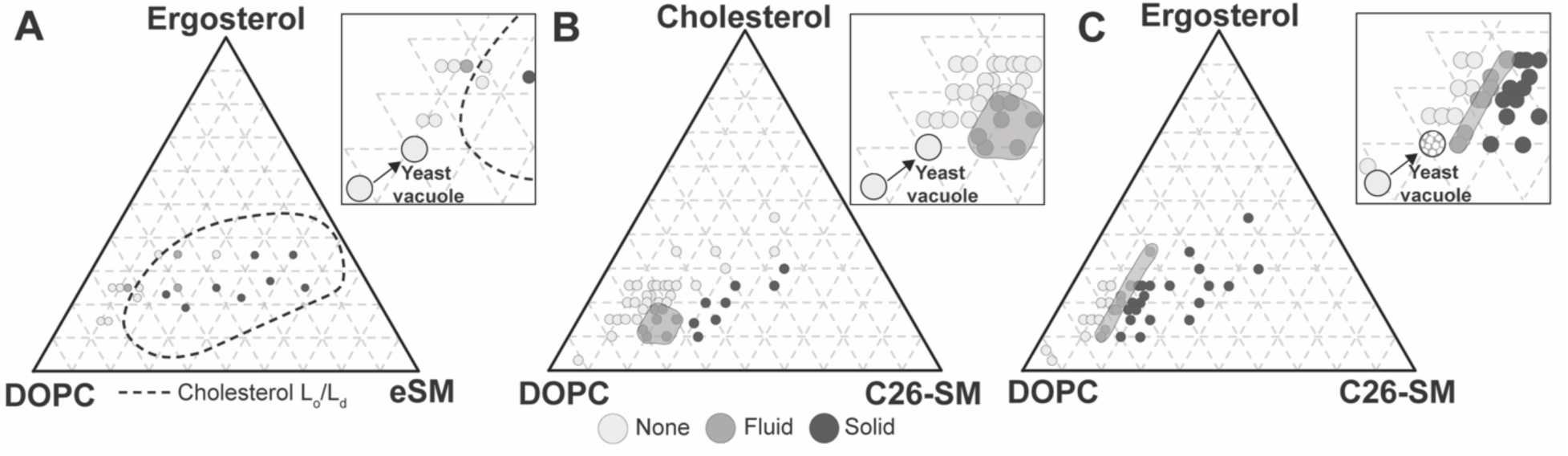
Ergosterol and cholesterol differentially support domain formation depending on sphingomyelin chain length. Shown are three experimentally sampled ternary phase diagrams of DOPC, one of two sphingomyelins, and one of two sterols constructed from sampling a range of GUV compositions via wide-field fluorescence microscopy. For each, points represent individually tested lipid compositions, colored by observed membrane behavior as determined by fluorescence microscopy. The shaded regions indicate the observed boundaries of regions in the phase diagram exhibiting fluid phase separation, reflecting likely *L*_o_/*L*_d_ coexistence. Insets for each panel show the approximate changes in composition of the yeast vacuole as it phase separates, plotted onto each corresponding phase map. (**A**) DOPC/eSM/ergosterol. The dashed outline indicates the reported *L*_o_/*L*_d_ coexistence region for DOPC/eSM/Cholesterol from prior literature, shown for comparison (39). (**B**) DOPC/C26-SM/cholesterol. (**C**) DOPC/C26-SM/ergosterol.

Replacing eSM with very long-chain C26-SM substantially shifted phase behavior of the systems tested. In cholesterol-containing membranes (Fig. 3B, Fig. S4), most compositions remained homogeneous, and liquid-liquid coexistence was restricted to a narrow region with intermediate C26-SM and cholesterol fractions. At higher C26-SM levels, membranes transitioned to those featuring solid domains. Ergosterol-containing membranes displayed a distinct arrangement of phases in the presence of C26-SM (Fig. 3C, Fig. S5). Liquid-liquid coexistence occurred within a narrow compositional window extending toward higher ergosterol fractions while C26-SM remained at moderate levels. This coexistence region lay between a homogeneous fluid regime at lower sphingomyelin fractions and an expanded solid-like region at higher C26-SM levels. While both C26-SM systems displayed a smaller region of liquid-liquid coexistence compared to eSM/cholesterol, these corresponded to lower sphingolipid and sterol concentrations that could be biologically relevant in the vacuole.

When yeast cells transition from late-exponential into early-stationary phase, their vacuole lipid composition changes through increases in ergosterol (7.0 ± 0.2 to 10.3 ± 0.1 mol% of polar lipids) and sphingolipids (3.6 ± 0.1 to 11.44 ± 1.0 mol% of polar lipids). This change corresponds to an increase in the fraction of vacuoles that are phase-separated from 0 to 80% of cells (21). We approximated these into a ternary composition change by considering phospholipid levels as equivalent to the low melting-temperature DOPC, given that most cellular phospholipids are unsaturated, and the total sphingolipids as level of the sphingomyelin. We used this simplification to ask if each of the analyzed systems approximated the behavior of the vacuole membrane. In the ergosterol/C26-SM/DOPC phase diagram (Fig. 3C, inset), exponential-phase vacuoles lie squarely within the homogeneous region, but shift close to the liquid-liquid coexistence regime during early stationary phase. In contrast, for both the eSM/ergosterol and eSM/cholesterol systems, the stationary-phase vacuole model remains far from the boundary into a phase-separated state. Thus, a combination of a long chain length and ergosterol in these simplified systems best mimics the phase behavior of the yeast vacuole membrane.

### Sterol-sphingolipid interactions alter membrane ordering in binary and ternary mixtures

The phase diagrams suggested that sterol structure and sphingomyelin chain length can each influence the formation of fluid membrane domains. To examine how these interactions affect membrane ordering that could underlie phase behavior, we measured Generalized Polarization of the solvatochromic dye Laurdan in large unilamellar vesicles (LUVs) (45, 46). Almost all sterols promote lipid ordering by condensing sphingolipid or phospholipid acyl chains (7, 12), but differential effects can be observed that reflect varying sterol-chain interactions (13). In binary sphingomyelin–sterol mixtures (80 mol % sphingomyelin and 20 mol % sterol), sterol structure strongly influenced membrane order when membranes contained egg sphingomyelin (eSM). Cholesterol-containing membranes produced higher generalized polarization (GP) values than ergosterol-containing eSM membranes across the temperature range examined (Fig. 4A), indicating increased eSM packing in the presence of cholesterol. A stronger condensing effect for cholesterol has been previously observed in other binary mixtures (47) and could underlie this sterol’s capacity to induce ordered membrane domain formation across a broader range of compositions (48).

**Figure 4.**
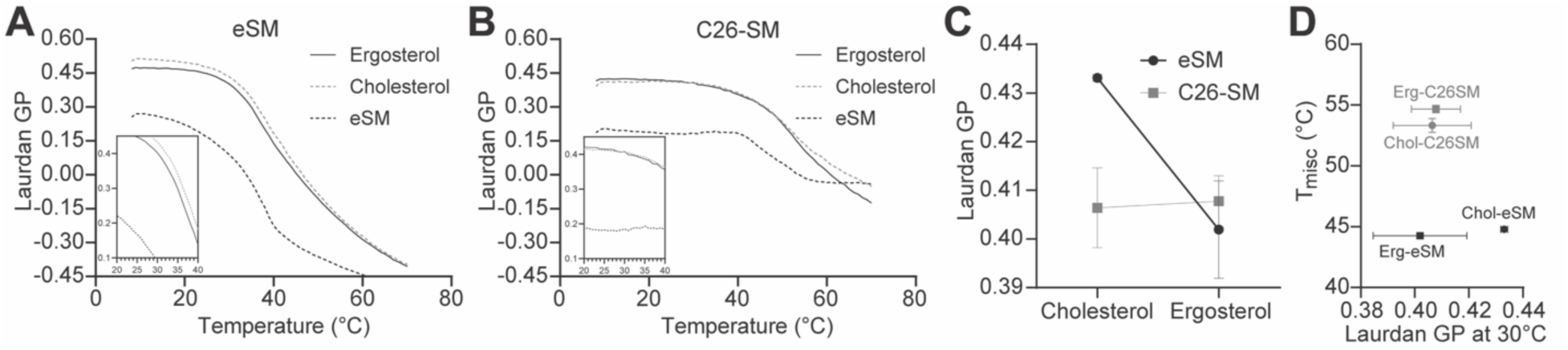
Sphingolipid chain length modulates sterol-dependent membrane order and transition temperatures in binary mixtures. (**A**) Laurdan generalized polarization (GP) as a function of temperature for 80/20 eSM/sterol mixtures containing cholesterol or ergosterol. Membranes display sterol-dependent differences in membrane order, with cholesterol systems showing a higher ordering and T_misc_. (**B**) Laurdan GP as a function of temperature for 80/20 C26-SM mixtures containing cholesterol or ergosterol. The C26-SM membranes exhibit elevated transition midpoints relative to eSM but eliminated T_misc_ differences between cholesterol and ergosterol. (**C**) Interaction between sphingomyelin and sterols for Laurdan GP at 30 °C, the optimal growth temperature for yeast. In eSM membranes, ergosterol lowers GP relative to cholesterol whereas C26-SM membranes show minimal sterol-dependent packing change at this temperature. (**D**) Laurdan GP at 30°C plotted against the transition midpoint (T_misc_) across compositions. The C26-SM systems show an elevated T_misc_, but a lack of Laurdan GP difference.

Like for phase properties, membranes containing C26-SM displayed different ordering parameters. GP values were elevated overall relative to eSM mixtures, reflecting the longer chain’s role in ordering. Similarly, transition midpoints derived from temperature-dependent GP curves were higher in C26-SM membranes than in eSM membranes (Fig. S6), consistent with the increased miscibility temperatures (T_misc_) for very-long-chain sphingolipids. However, sterol-dependent differences in both ordering and T_misc_ were substantially reduced compared to eSM (Fig. 4B). This interaction is illustrated by comparing GP values at 30 °C, a common yeast growth temperature. Here, ergosterol lowers GP relative to cholesterol in eSM membranes, but produces minimal changes in membranes containing C26-SM (Fig. 4C). C26-SM eliminates the influence of sterol structure on membrane ordering, despite its overall capacity to increase T_misc_ compared to ergosterol (Fig. 4D).

To further assess how temperature, sphingolipid chain length and sterol structure interact to influence membrane order, we performed a three-way ANOVA on Laurdan GP values from 80/20 sphingomyelin/sterol binary mixtures (Table 1, Table S1). Temperature had a strong effect on GP (p < 0.0001) with higher temperatures decreasing GP. Chain length showed a borderline effect (p = 0.051) while sterol structure had a moderate effect (p = 0.026), with ergosterol producing lower GP values than cholesterol. Notably, there was significant interaction between temperature and sphingolipid chain length (p < 0.0001), suggesting very long chain sphingolipids (C26-SM) are less sensitive to temperature-induced disordering compared to eSM. In contrast, interactions between sterol and sphingolipid (p = 0.088) or temperature and sterol (p = 0.447) were not significant and no three-way interaction was observed (p = 0.872). Together, this indicates that GP is influenced by temperature and that very long chain sphingolipids stabilize membrane packing and reduce thermal sensitivity.

**Table 1.** Three-factor ANOVA evaluating the effects of temperature, sphingolipid chain length (EggSM vs C26-SM) and sterol structure (cholesterol vs ergosterol) along with their interactions on the measured response variable. P-values for each term are shown. The overall model was significant (F(7,16) =61.19, p < 0.0001) with R^2^ = 0.964.

| Source of variation | p value |
| --- | --- |
| Temperature | <0.0001 |
| Sphingolipid chain length | 0.051 |
| Sterol | 0.026 |
| Temperature x SM | <0.0001 |
| Temperature x Sterol | 0.447 |
| SM x Sterol | 0.088 |
| Temperature x SM x Sterol | 0.872 |

We next asked whether these sterol–sphingolipid interactions persist in more complex membranes used for GUV experiments (Fig. 5). In ternary mixtures containing DOPC, C26-SM, and sterol, we observed that cholesterol systems showed enhanced ordering compared to ergosterol systems at low C26-SM and high DOPC levels (Fig. 5A). Previous studies have shown that ergosterol has poor condensing effects on unsaturated PC species, explaining this difference (49). However, as C26-SM levels rose, the difference between the ergosterol and cholesterol systems was eliminated (Fig. 5B-C), before recovering somewhat when both sphingomyelin and sterol levels were both at high concentrations (Fig. 5D). The latter effect is consistent with observations that cholesterol continues ordering membranes even at high concentrations, while ergosterol loses efficacy (49, 50).

**Figure 5.**
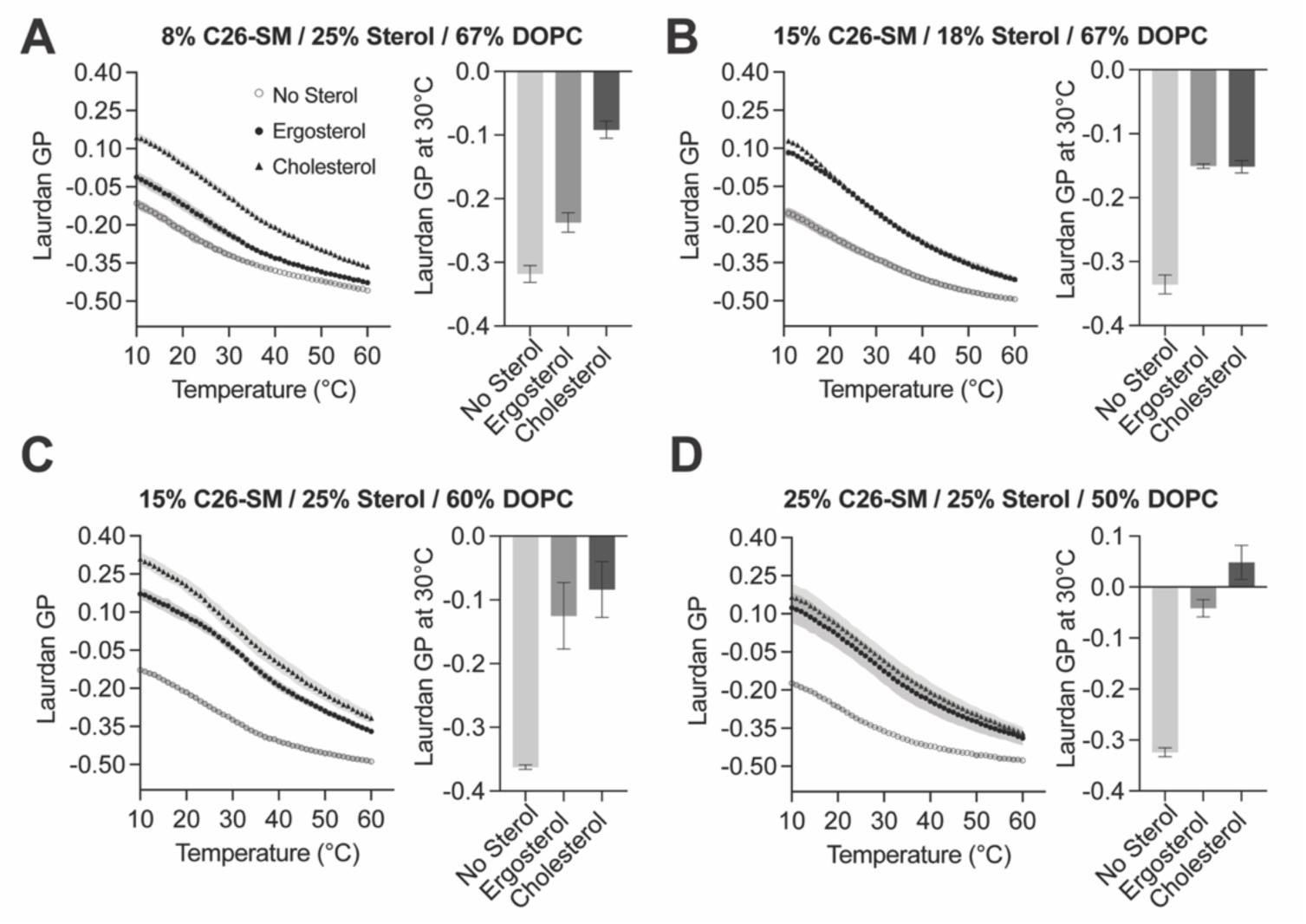
Variations of membrane order in C26-SM-containing ternary mixtures. Each panel shows Laurdan GP of LUVs as a function of temperature for selected liposome compositions spanning varying C26-SM and sterol species. Compositions were chosen to highlight conditions where sterol-dependent differences in Laurdan GP are either pronounced or minimal. (**A**) In low C26-SM and high sterol membranes, cholesterol-containing membranes show higher ordering (GP) than those with ergosterol, which shows only a small difference from sterol-free membranes. When C26-SM is increased to 15 mol %, similar to its sphingolipid concentration in the yeast vacuole, ordering differences between ergosterol and cholesterol disappear for both 18 (**B**) and 25 (**C**) mol % sterol systems. At high C26-SM and high sterol levels (**D**), cholesterol regains a modest ordering capacity, which could be related to the higher solubility of cholesterol in membranes compared to ergosterol (57).

Together, these measurements indicate that sterol-dependent differences in membrane order observed in membranes containing shorter sphingolipids become attenuated when sphingolipid acyl chains are very long. This shift parallels the changes in phase behavior observed in the GUV phase diagrams, where cholesterol-containing membranes more readily support fluid domain formation in eSM mixtures but not in membranes containing C26 sphingomyelin. In contrast, ergosterol-containing membranes retain a compositional window that supports fluid phase coexistence in the presence of very-long-chain sphingolipids.

## Discussion

Membrane phase separation into coexisting liquid domains depends on the cooperative interactions between sterols and saturated lipids like sphingolipids. Both these lipid classes differ substantially across eukaryotic cell types. Understanding if changes in one lipid class are relevant to its interaction with the other provides insights into the evolution of eukaryotic lipid metabolism, as well as the underlying mechanism for membrane phase behavior. Here we examined how one aspect of sphingolipid structure – the length of its *N*-acyl chain length – interacts with two sterol classes, ergosterol and cholesterol. The motivation lay in the coincidence of very long chain sphingolipids in fungal cells alongside ergosterol, while mammalian plasma membranes contain shorter sphingolipids alongside cholesterol. In yeast, the organization of the vacuolar membrane into micron-scale domains as cells enter stationary phase was also lost when very long-chain sphingolipid synthesis was lost, or when ergosterol was replaced with cholesterol through genetic engineering. Although we cannot rule out that trafficking of sphingolipids and/or sterols to the vacuole was impeded by these perturbations, they suggested that structural changes in both molecules could influence domain formation.

To investigate the physical basis of lipid-dependencies in the vacuole, we reconstructed sterol–sphingolipid membranes in GUVs and mapped their phase behavior across composition space. In membranes containing C16-chain eSM, ergosterol did not consistently support liquid–liquid coexistence, in contrast to previously reported phase diagrams of cholesterol-containing membranes with eSM (38, 39). In C26-SM membranes, ergosterol and cholesterol both showed capacities to support liquid-liquid coexistence albeit at different stoichiometries. Increasing sphingolipid chain length is known to increase bilayer thickness and alter packing interactions between neighboring lipids (16). Very long acyl chains can extend across the bilayer midplane and promote interdigitation between opposing leaflets (17, 51), while chain length can generate hydrophobic mismatch between lipid species that influences lateral organization in membranes (52, 53). These constraints likely reduce the ability of cholesterol to accommodate very long sphingolipid chains within fluid ordered domains and could explain why C26-SM inhibits fluid domain formation compared to eSM for cholesterol systems. In contrast, ergosterol’s more moderate ordering capacity could restrict phase separation of eSM-containing systems.

Importantly, liquid domains could be generated for lower sterol and sphingolipid concentrations for the C26-SM systems than for eSM. For ergosterol/C26-SM/DOPC, this reduced threshold corresponded to previous measurements for the concentrations of these components – approximately 12 mol% sphingolipids and 10 mol% ergosterol – in purified vacuoles that are phase-separated (21). These sterol and sphingolipid concentrations are rather modest compared to commonly used model membrane compositions, or those found in the mammalian plasma membrane (54). Membrane domains in the yeast vacuole also display a melting temperature that is actively tuned by growth temperature, implying that its lipidome in stationary-phase cells is maintained in a state near a phase boundary. This phenomenon is also consistent with the C26-SM/ergosterol/DOPC phase diagram, which shows that vacuole-levels of ergosterol and sphingolipids are found near the phase boundary between homogenous and domain-containing membranes. This suggests that modest changes in both sphingolipid and ergosterol levels could be used to induce the vacuole membrane to enter or leave a two-phase regime, as has been measured changes in phospholipids that alter saturation of their acyl chains (31). Thus, even though C26-SM is not a fungal sphingolipid due to its headgroup and lack of hydroxylations, its fungal-like C26 *N*-acyl chain induces ternary mixtures to behave more closely to what is observed in cell membranes in several respects.

To further explore the basis for GUV phase behavior, we measured the ordering interactions between sphingomyelin chain length and sterol identity in bulk LUVs using Laurdan fluorescence spectroscopy. Differences between the ability of ergosterol and cholesterol to promote chain ordering have been widely reported, reflecting how sterol ring structure and side-chain geometry influence packing interactions with saturated lipid chains (8, 11, 12, 14). We confirmed previous observations that ergosterol induced more moderate ordering of saturated long-chain lipids, like eSM, compared to cholesterol. This effect could be relevant for the poor capacity of ergosterol/eSM mixtures to phase-separate, since formation of ordered domains is thought to require increased packing differences between the saturated lipid-enriched *L*_o_ phase and the unsaturated lipid-enriched *L*_d_ phase. Importantly, we find that the difference between ergosterol and cholesterol is lost for C26-SM. C26-SM, like glucosylceramide (34), predominantly forms solid domains in cholesterol and ergosterol mixtures, consistent with the high T_misc_ of C26-SM/sterol mixtures. However, both cholesterol and ergosterol systems show a distinct region of fluid domain coexistence. There are some key differences in the concentration regimes in which the two sterols support this with C26-SM. The ergosterol systems form fluid domains at generally lower levels of C26-SM/sterol, which are more relevant to the lipidomes measured in the yeast vacuole. Ternary mixtures reflecting these concentrations show identical ordering for ergosterol and cholesterol, but those with reduced C26-SM show greater ordering for cholesterol (Fig. 5A). The ergosterol/C26-SM systems also show an elevated T_misc_ compared to their ordering, which could contribute to phase separation (Fig. 5B). Overall, while some of the ordering differences between the systems are subtle, they translate into significant phase property differences that could be biologically relevant.

## Conclusions

Membrane phase separation depends on the cooperative interactions between sterols and sphingolipids. We examined how sterol structure and sphingolipid acyl chain length together influence membrane organization using a combination of yeast genetics and reconstituted model membranes. Vacuolar membranes in wild-type yeast form micron-scale domains during stationary phase, whereas disruption of very-long-chain sphingolipid synthesis or replacement of ergosterol with cholesterol impaired this organization in vivo. To determine the physical basis of these effects, we reconstructed sterol–sphingolipid membranes in giant unilamellar vesicles and mapped their phase behavior across composition space. Small changes in lipid structure produced large changes in membrane phase behavior. In membranes containing C16-chain eSM, ergosterol produced liquid–liquid coexistence only within a relatively narrow region of the phase diagram. By comparison, previously reported phase diagrams of cholesterol-containing membranes with eSM display liquid–liquid coexistence across a broader range of compositions (38, 39). In contrast, membranes containing C26-SM displayed a different pattern: cholesterol mixtures were largely homogeneous across most compositions, while ergosterol mixtures supported liquid–liquid coexistence within a distinct compositional window positioned between homogeneous membranes and solid-like phases. These results suggest that sterol and sphingolipid structures may be functionally matched in biological membranes, linking sterol chemistry with the distinctive lipid composition of fungal membranes. More broadly, this relationship highlights how small variations in lipid structure can reorganize membrane phase behavior and suggests that co-evolution of sterols and sphingolipids may help tune membrane physical properties across eukaryotic lineages.

## Supporting information

Combined Supplementary Information

## Acknowledgements

Research was supported by the National Institute of General Medical Sciences (R35-GM142960 to I.B., T32-GM008326C to I.J-C.), National Science Foundation (MCB-2046303 to I.B.), and Department of Energy (DE-SC0022954 to I.B.).

## Author contributions

I.J-C. and I.B. conceived the project and designed the experiments. I.J-C. performed the experiments and analyzed the data. H.K. generated strains and performed analysis. I. J-C. and I.B. wrote the manuscript.

## Materials and Methods

### Strain generation

*Saccharomyces cerevisiae* strains used in this study were derived from the W303a background. Gene deletions were generated by PCR-based homologous recombination using standard lithium acetate transformation methods. Deletion of *ELO3* was performed and replaced with a *KANMX* resistance cassette. Similarly, deletion of *ELO2* was achieved using a *LEU2* selection marker. The strain engineering of the cholesterol yeast was performed as described elsewhere (32). Briefly, yeast codon-optimized *Danio rerio* DHCR7 and DHCR24 gene fragments were synthesized (VectorBuilder) and integrated into the yeast genome by homologous recombination. *DHCR7* and *DHCR24* were inserted in place of the endogenous ERG6 and ERG5 loci, respectively. Both genes were expressed under the control of the constitutive TDH3 promoter.

### Cell growth and vacuole domain analysis

Cells were grown in yeast extract–peptone–dextrose (YPD) medium or in synthetic complete medium containing 0.5% ammonium sulfate, 0.17% yeast nitrogen base without amino acids, and 2% glucose. For microscopy experiments examining vacuole membrane domains, cells were grown to stationary phase under glucose-depleted conditions to induce vacuole membrane domain formation as described previously (55). Briefly, a single colony was initially inoculated in rich medium (YPD) and grown overnight, then diluted to synthetic complete medium with 2% glucose and allowed to continue growth before being diluted into minimal medium containing reduced glucose (0.4%) supplemented with essential amino acids and nucleotide bases. Cells were imaged after immobilization on concanavalin A coated eight-well chambered coverglasses (1-2 mg/mL) to facilitate adherence during acquisition.

Vacuoles were visualized at room temperature using a Zeiss LSM 880 Airyscan equipped with a Plan-Apochromat 63x/1.4 NA oil immersion objective and an Airyscan detector. Pho-GFP was excited with a 488 nm laser line at 2% power. Z-stacks spanning the top and bottom of the vacuole were acquired and processed using default Airyscan settings. 3D projections of Z-stacks were generated in ImageJ for downstream analysis. Vacuoles within large fields of cells were scored manually and categorized. Briefly, vacuoles were classified as uniform (continuous, regularly patterned domains), or homogenous (no detectable domain formation within resolution limits of Airyscan imagining). For each biological replicate, at least 100 cells were analyzed.

### Sterol extraction and analysis

Sterol extraction from yeast was performed as described previously with minor modifications (56). Briefly, cells corresponding to an optical density of 1 at OD₆₀₀ were supplemented with an internal standard and resuspended in 1 mL of Milli-Q water. Cholesterol was used as the internal standard for wild-type (ergosterol-producing) strains, whereas ergosterol was used for cholesterol-producing strains. Cells were lysed by bead beating using an equal volume of zirconia beads (BioSpec), and the lysate was transferred to a glass tube containing a Teflon-lined cap. Saponification was carried out by adding 3 mL of methanolic KOH and incubating the mixture at 70 °C for 2 h.

Following cooling to room temperature, sterols were extracted by addition of 3 mL of *n*-hexane, vortexing briefly, and centrifugation (5 min, 800 rpm). The upper organic phase was collected, and the aqueous phase was subjected to a second extraction. Combined organic fractions were dried under a stream of nitrogen. Dried extracts were derivatized in a 1:1 mixture of pyridine and trimethylchlorosilane at 70 °C for 1 h prior to analysis. Samples were analyzed by gas chromatography–mass spectrometry (GC–MS) using an Agilent 8890 GC coupled to a 5977B mass spectrometer equipped with a 30 m DB-5 column. The oven temperature was programmed as follows: 120 °C (1 min hold), ramp to 270 °C at 20 °C/min, followed by a slower ramp to 290 °C at 2 °C/min, and a final ramp to 300 °C at 20 °C/min with a 2 min hold.

Sterol identification was confirmed by comparison to the NIST17 mass spectral library. Quantification was performed using calibration curves generated from external standards, with values corrected for extraction efficiency based on recovery of the internal standard in each sample

### Reagents

1,2-dipalmitoyl-sn-glycero-3-phosphocholine (DOPC), and Egg Sphingomyelin (eSM) were obtained from Avanti Polar Lipids. Ergosterol was obtained from Thermo Fisher Scientific. Cholesterol and C26:0 sphingomyelin (C26-SM) were obtained from Cayman Chemical. The fluorescent *L*_d_ marker Texas Red DHPE was obtained from Life Technologies. All lipids were used without further purification. Cholesterol and ergosterol were mixed from powder in chloroform to a concentration of 10 mg/mL. Texas Red DHPE was mixed in chloroform to a concentration of 0.5 mg/mL. C26-SM was mixed in chloroform to a concentration of 2 mg/mL. Lipids obtained from Avanti Polar Lipids were supplied in chloroform at nominal concentrations of 10 mg/mL and were used without further verification. All other chemical reagents were from Sigma-Alrdich.

### Electroformation of GUVs

Vesicles composed of DOPC, sphingomyelin, and sterol mixtures were generated by electroformation. Glass slides coated with indium tin oxide (ITO; Delta Technologies) were heated on a heat block at 60 °C. Lipid stock solutions in chloroform containing 0.8 mol% Texas Red–DHPE were mixed to achieve a total of 0.25 mg lipid and briefly warmed to 60 °C. The lipid mixture was then spread evenly onto the ITO-coated slides. After solvent evaporation, the slides were placed under vacuum for at least 30 min to remove residual chloroform. The two slide halves were assembled face-to-face and separated by a Viton O-ring (Ace Glass, 7855-14), and the chamber was filled with 200 mM sucrose solution. An AC voltage of 1.3 V at 10 Hz was applied across the chamber for 1 h at 75 °C to induce vesicle formation. The electroformation temperature was chosen to exceed the highest lipid transition temperature present in the mixtures. Following electroformation, the chamber was transferred to a 60 °C oven for at least 30 min, then to 37 °C for at least an additional 30 min and finally equilibrated at room temperature for 1 h to allow gradual cooling of the membranes and promote nucleation and coalescence of membrane domains.

### Imaging of GUVs

GUV suspensions were diluted in 200 mM glucose solution and imaged using wide-field fluorescence microscopy (Thermo Fisher EVOS equipped with a 63× Nikon oil-immersion objective). For each preparation, at least 30 GUVs were analyzed and manually classified based on domain morphology following the 1 h equilibration period at room temperature described above. Vesicles containing a single coalesced domain were classified as phase-separated. Vesicles displaying multiple discrete domains that remained after equilibration, or domains with irregular morphologies, were classified as solid-like or gel-like. Vesicles exhibiting a uniform distribution of Texas Red–DHPE were classified as having no detectable domains. Images were processed in ImageJ/Fiji.

### LUV preparation

Large unilamellar vesicles with defined lipid compositions were prepared by lipid extrusion. Lipids dissolved in chloroform were mixed to achieve the desired molar ratios and dried under a stream of nitrogen to form a thin lipid film. Residual solvent was removed under vacuum for at least 1 h. The dried lipids were rehydrated in 20 mM HEPES pH 7.5 buffer, and vesicles were extruded 21 times through 100 nm polycarbonate membranes using an extruder pre-equilibrated to 70 °C.

### Laurdan GP analysis

Steady-state fluorescence measurements were performed on a Cary Eclipse fluorometer (Agilent) equipped with automated polarizers and temperature control. Liposomes prepared as described above were stained with Laurdan (Invitrogen D250) from a 5 µM stock solution at a 1:100 dilution. Samples were excited at 365 nm and emission intensities were recorded at 440 nm and 490 nm. Generalized polarization (GP) values were calculated as a unitless ratio based on emission intensities at the two wavelengths:

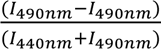

GP values were measured during temperature sweeps from 8 °C to 60-70 °C at a heating rate of 1 °C min^-1^.

### Determination of GP transition midpoints

Laurdan GP values obtained during temperature sweeps were fit to a sigmoidal Boltzmann function using nonlinear least-squares regression in GraphPad Prism (GraphPad Software). The Boltzmann equation was defined as:

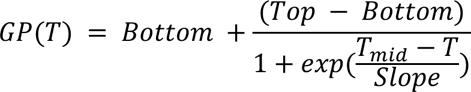

where Bottom and Top represent the lower and upper GP plateaus, T is temperature, T_mid_ is the midpoint of the transition, and Slope describes the steepness of the transition. The transition midpoint T_mid_ was extracted from the fitted curves for each lipid composition tested and was considered as the T_misc_ for the lipid mixture.

### Statistical analyses

Statistical analyses were performed using GraphPad Prism (GraphPad Software). Linear regression was used to examine relationships between GP values and transition midpoints across lipid compositions. To evaluate the effects of sterol species, sphingomyelin chain length, and temperature, multiple linear regression models including interaction terms were fit using least-squares estimation. Model coefficients and associated statistical significance were determined within Prism.

