## Supplementary material for "Coupling between sterol and sphingolipid structure in ordered membrane domains": Combined Supplementary Information

This file contains:

Figure S1-S6

Table S1-S2

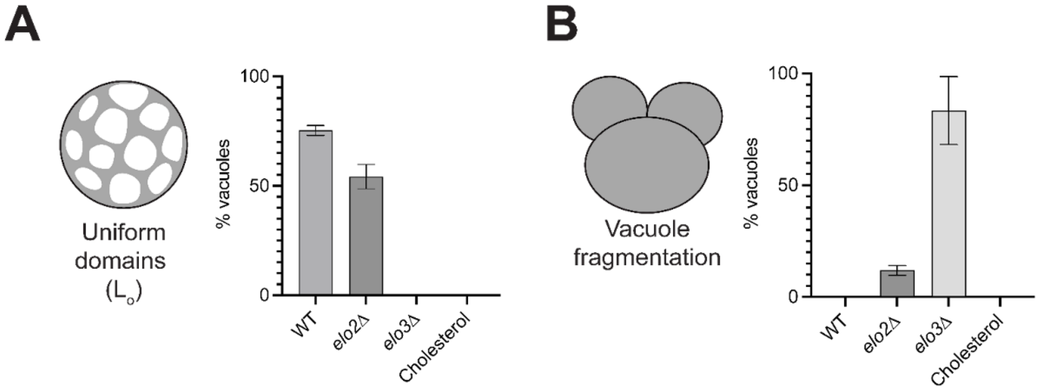


**Figure S1. Sphingolipid chain length and sterol composition influence vacuolar domain formation and fragmentation.** (**A**) Percentage of vacuoles exhibiting visible liquid-ordered (Lo) membrane domains. (**B**) Percentage of vacuoles exhibiting vacuole fragmentation. Cartoons illustrate the phenotypes scored in each analysis. Values represent mean +/- SD from three biological replicates with > 100 vacuoles quantified per replicate.

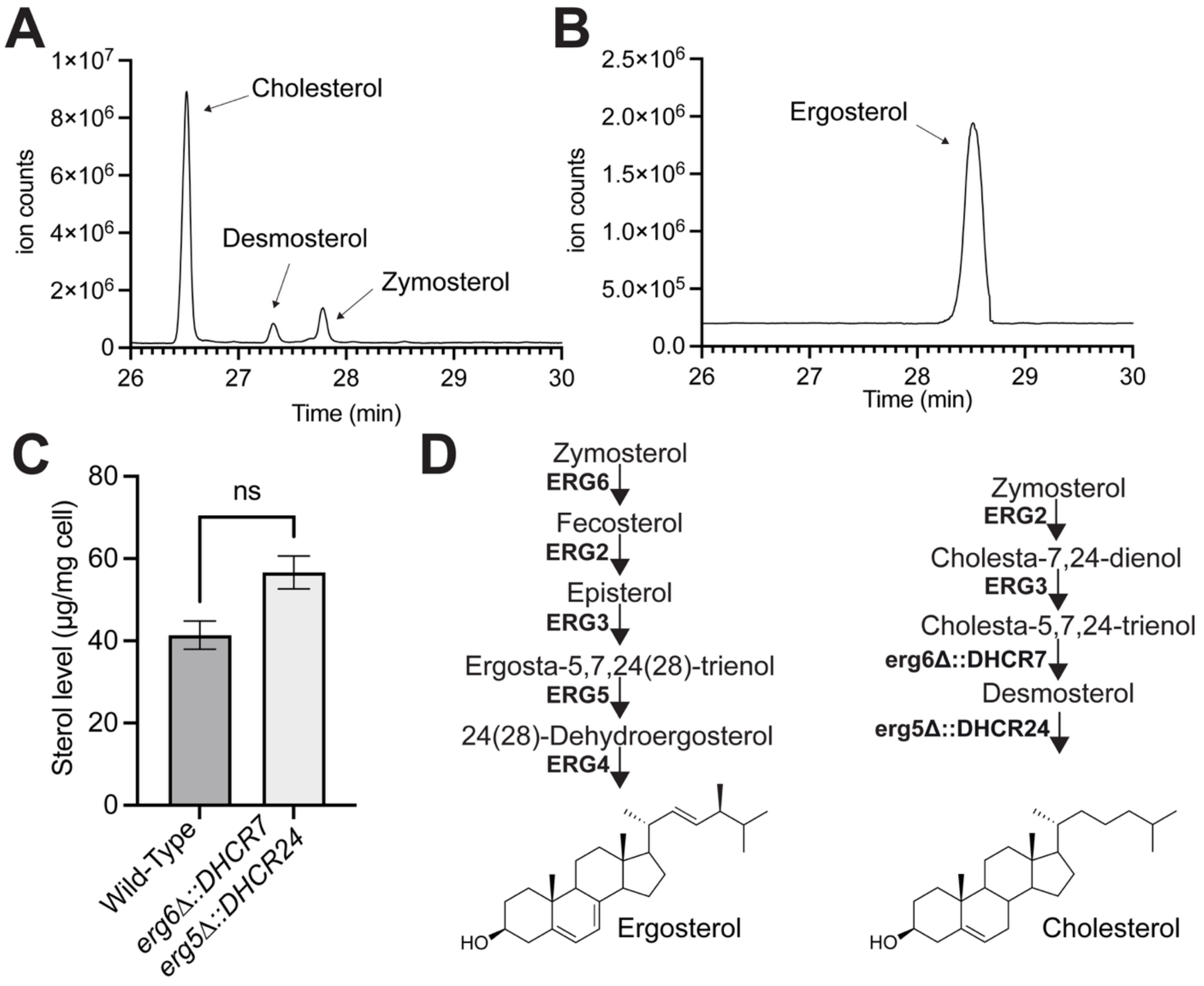


**Figure S2. GC-MS validation of cholesterol producing strain.** (**A**) Representative GC chromatogram of sterols extracted from engineered CHOL yeast. Peaks corresponding to cholesterol, desmosterol, and zymosterol are indicated. Compound identities were confirmed by comparison of the associated mass spectra to entries in the NIST mass spectral library. (**B**) Identical analysis of WT yeast synthesizing ergosterol. (**C**) Quantification of sterol abundance from GC-MS analysis showing cholesterol as the predominant sterol species. Bars represent mean +/- SD from three biological replicates. (**D**) Pathway for synthesizing ergosterol in WT yeast (left) and cholesterol in the CHOL strain (right).

**A**

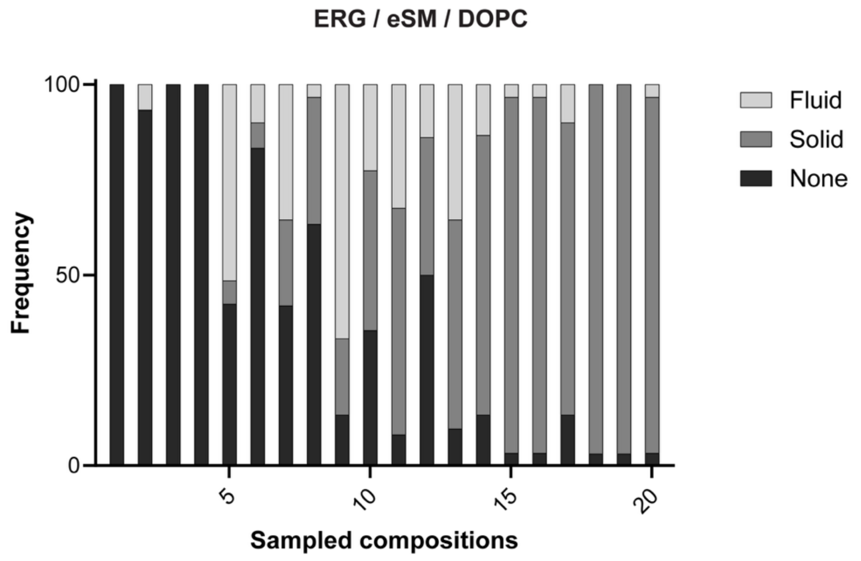


**B**

| Comp | eSM (mol%) | Ergosterol (mol%) | DOPC (mol%) | n | Comp | eSM (mol%) | Ergosterol (mol%) | DOPC (mol%) | n |
|---|---|---|---|---|---|---|---|---|---|
| 1 | 8 | 25 | 67 | 30 | 11 | 30 | 19 | 51 | 37 |
| 2 | 10 | 15 | 75 | 30 | 12 | 30 | 35 | 35 | 36 |
| 3 | 10 | 25 | 65 | 30 | 13 | 35 | 25 | 40 | 31 |
| 4 | 12 | 15 | 73 | 32 | 14 | 40 | 35 | 25 | 30 |
| 5 | 12 | 25 | 63 | 33 | 15 | 43 | 22 | 35 | 30 |
| 6 | 15 | 25 | 60 | 30 | 16 | 48 | 27 | 25 | 30 |
| 7 | 15 | 35 | 50 | 31 | 17 | 50 | 35 | 15 | 30 |
| 8 | 16 | 22 | 62 | 30 | 18 | 53 | 25 | 22 | 32 |
| 9 | 20 | 35 | 45 | 30 | 19 | 55 | 35 | 10 | 32 |
| 10 | 23 | 23 | 54 | 31 | 20 | 58 | 26 | 16 | 30 |

**Figure S3. Classification of phase behavior across sampled ERG/eSM/DOPC compositions.** (**A**) Frequency of vesicles displaying different phase behaviors for each sampled lipid composition. Vesicles were classified as fluid, solid or none based on Texas Red DPHE partitioning as described in methods. Bars represent the percent of the vesicles assigned to each category for a given composition. The x-axis represents the sampled compositions. (**B**) Lipid compositions corresponding to each sampled composition shown in panel A. Each composition is defined by the molar fraction of eSM, ergosterol, and DOPC. The column labeled Comp corresponds to the compositions plotted to align the x-axis in panel A. The column n indicates the number of vesicles analyzed for that composition.

A

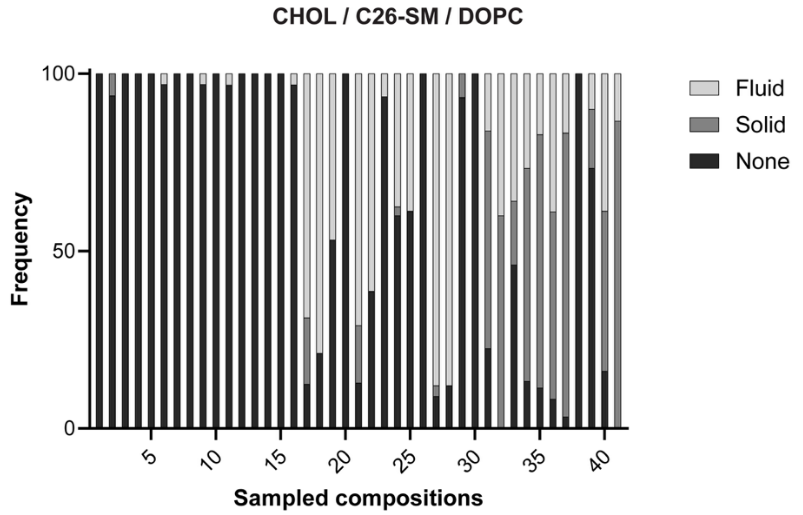


B

| Comp | C26-SM (mol%) | Cholesterol (mol%) | DOPC (mol%) | n |
|---|---|---|---|---|
| 1 | 8 | 15 | 77 | 31 |
| 2 | 8 | 25 | 67 | 32 |
| 3 | 10 | 15 | 75 | 35 |
| 4 | 10 | 20 | 70 | 33 |
| 5 | 10 | 25 | 65 | 30 |
| 6 | 12 | 15 | 73 | 33 |
| 7 | 12 | 20 | 68 | 32 |
| 8 | 14 | 22 | 64 | 32 |
| 9 | 14 | 25 | 61 | 33 |
| 10 | 15 | 15 | 70 | 30 |
| 11 | 15 | 18 | 67 | 31 |
| 12 | 15 | 20 | 65 | 32 |
| 13 | 15 | 22 | 63 | 35 |
| 14 | 15 | 25 | 60 | 30 |
| 15 | 15 | 35 | 50 | 33 |

| Comp | C26-SM (mol%) | Cholesterol (mol%) | DOPC (mol%) | n |
|---|---|---|---|---|
| 16 | 17 | 25 | 58 | 32 |
| 17 | 18 | 12 | 70 | 32 |
| 18 | 18 | 18 | 64 | 33 |
| 19 | 18 | 20 | 62 | 32 |
| 20 | 18 | 25 | 57 | 34 |
| 21 | 20 | 10 | 70 | 31 |
| 22 | 20 | 15 | 65 | 31 |
| 23 | 20 | 18 | 62 | 31 |
| 24 | 20 | 20 | 60 | 40 |
| 25 | 20 | 22 | 58 | 31 |
| 26 | 20 | 25 | 55 | 30 |
| 27 | 25 | 10 | 65 | 33 |
| 28 | 25 | 15 | 60 | 33 |
| 29 | 25 | 25 | 50 | 30 |
| 30 | 25 | 35 | 40 | 31 |

| Comp | C26-SM (mol%) | Cholesterol (mol%) | DOPC (mol%) | n |
|---|---|---|---|---|
| 31 | 30 | 14 | 56 | 31 |
| 32 | 30 | 20 | 50 | 30 |
| 33 | 30 | 30 | 40 | 39 |
| 34 | 33 | 10 | 57 | 30 |
| 35 | 35 | 15 | 50 | 35 |
| 36 | 35 | 20 | 45 | 36 |
| 37 | 35 | 25 | 40 | 30 |
| 38 | 35 | 45 | 20 | 30 |
| 39 | 40 | 35 | 25 | 30 |
| 40 | 45 | 25 | 30 | 31 |
| 41 | 45 | 30 | 25 | 30 |

**Figure S4. Classification of phase behavior across sampled CHOL/C26-SM/DOPC compositions.** (**A**) Frequency of vesicles displaying different phase behaviors for each sampled lipid composition. Vesicles were classified as fluid, solid or none based on Texas Red DPHE partitioning as described in methods. Bars represent the percent of the vesicles assigned to each category for a given composition. The x-axis represents the sampled compositions. (**B**) Lipid compositions corresponding to each sampled composition shown in panel A. Each composition is defined by the molar fraction of C26-SM, cholesterol, and DOPC. The column labeled Comp corresponds to the compositions plotted to align the x-axis in panel A. The column n indicates the number of vesicles analyzed for that composition.

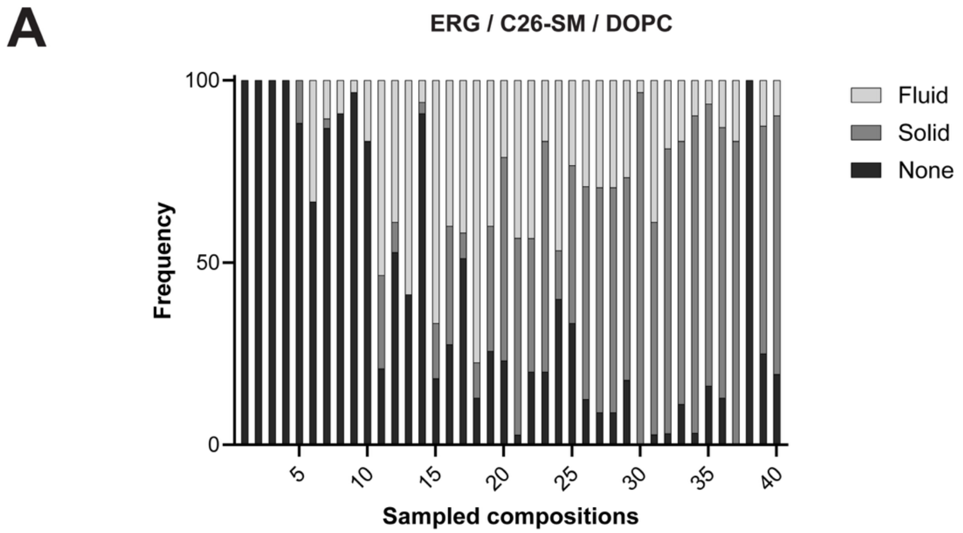


B

| Comp | C26-SM (mol%) | Ergosterol (mol%) | DOPC (mol%) | n |
|---|---|---|---|---|
| 1 | 8 | 15 | 77 | 33 |
| 2 | 8 | 25 | 67 | 32 |
| 3 | 10 | 15 | 75 | 42 |
| 4 | 10 | 20 | 70 | 30 |
| 5 | 10 | 25 | 65 | 34 |
| 6 | 11 | 10 | 79 | 33 |
| 7 | 12 | 15 | 73 | 38 |
| 8 | 12 | 20 | 68 | 33 |
| 9 | 14 | 18 | 68 | 30 |
| 10 | 14 | 22 | 64 | 30 |
| 11 | 15 | 10 | 75 | 43 |
| 12 | 15 | 12 | 73 | 36 |
| 13 | 15 | 18 | 67 | 34 |
| 14 | 15 | 20 | 65 | 33 |
| 15 | 15 | 25 | 60 | 33 |

| Comp | C26-SM (mol%) | Ergosterol (mol%) | DOPC (mol%) | n |
|---|---|---|---|---|
| 16 | 15 | 35 | 50 | 40 |
| 17 | 17 | 25 | 58 | 43 |
| 18 | 18 | 18 | 64 | 31 |
| 19 | 18 | 20 | 62 | 35 |
| 20 | 18 | 25 | 57 | 52 |
| 21 | 20 | 10 | 70 | 37 |
| 22 | 20 | 15 | 65 | 30 |
| 23 | 20 | 18 | 62 | 30 |
| 24 | 20 | 20 | 60 | 30 |
| 25 | 20 | 22 | 58 | 30 |
| 26 | 20 | 25 | 55 | 48 |
| 27 | 25 | 10 | 65 | 34 |
| 29 | 25 | 15 | 60 | 45 |
| 30 | 25 | 25 | 50 | 36 |

| Comp | C26-SM (mol%) | Ergosterol (mol%) | DOPC (mol%) | n |
|---|---|---|---|---|
| 31 | 25 | 35 | 40 | 32 |
| 32 | 30 | 30 | 40 | 36 |
| 33 | 35 | 15 | 50 | 31 |
| 34 | 35 | 20 | 45 | 31 |
| 35 | 35 | 25 | 40 | 31 |
| 36 | 35 | 45 | 20 | 30 |
| 37 | 3 | 6 | 91 | 30 |
| 38 | 40 | 25 | 35 | 32 |
| 39 | 45 | 30 | 25 | 31 |

**Figure S5. Classification of phase behavior across sampled ergosterol/C26-SM/DOPC compositions.** (**A**) Frequency of vesicles displaying different phase behaviors for each sampled lipid composition. Vesicles were classified as fluid, solid or none based on Texas Red DPHE partitioning as described in methods. Bars represent the percent of the vesicles assigned to each category for a given composition. The x-axis represents the sampled compositions. (**B**) Lipid compositions corresponding to each sampled composition shown in panel A. Each composition is defined by the molar fraction of C26-SM, ergosterol, and DOPC. The column labeled Comp corresponds to the compositions plotted to align the x-axis in panel A. The column n indicates the number of vesicles analyzed for that composition.

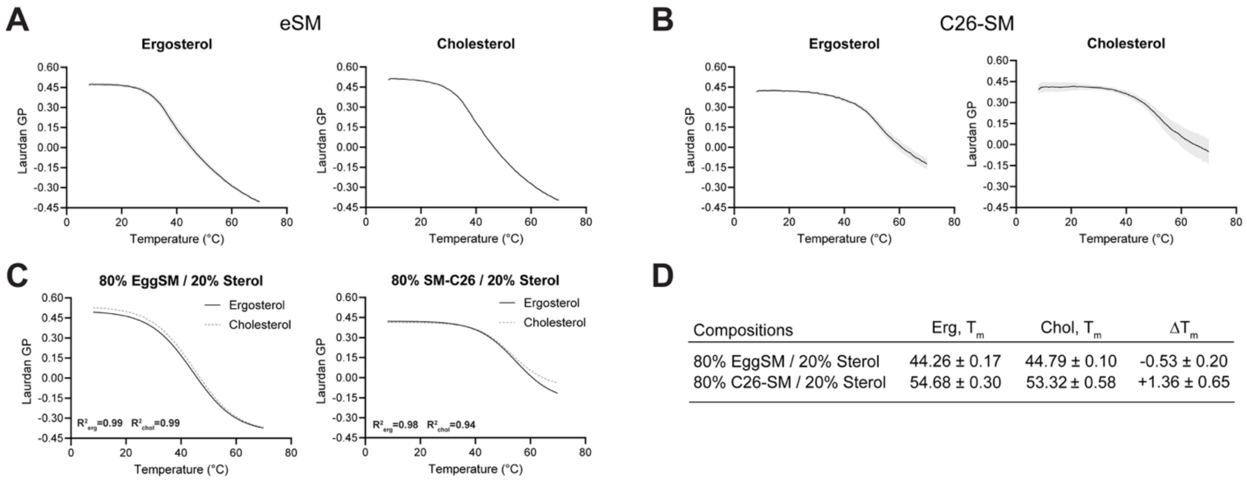


| Compositions | Erg, $T_m$ | Chol, $T_m$ | $\Delta T_m$ |
|---|---|---|---|
| 80% EggSM / 20% Sterol | 44.26 ± 0.17 | 44.79 ± 0.10 | -0.53 ± 0.20 |
| 80% C26-SM / 20% Sterol | 54.68 ± 0.30 | 53.32 ± 0.58 | +1.36 ± 0.65 |

**Figure S6. Membrane ordering in sterol/sphingomyelin binary systems.** (**A**) Laurdan generalized polarization (GP) as a function of temperature for bilayers composed of 80 mol% eSM and 20 mol% sterol, containing either ergosterol or cholesterol. Points represent the mean of three independent liposome preparations. (**B**) Laurdan GP temperature profiles for bilayers composed of 80 mol% C26-SM and 20 mol% sterol. (**C**) Boltzmann fits to the Laurdan GP curves used to determine the midpoint transition temperature ($T_m$). (D) $T_m$ values derived from Boltzmann fits for each composition. $\Delta T_m$ indicates the difference between ergosterol and cholesterol containing membranes (Erg - Chol).

| Source of Variation | β | 95% CI | p value | |
|---|---|---|---|---|
| ***Main Effects*** | | | | |
| Temperature | −0.154 | −0.181 – −0.127 | <0.0001 | **** |
| SM | −0.027 | −0.054 – 0.002 | 0.051 | ns |
| Sterol | −0.031 | −0.058 – −0.004 | 0.026 | * |
| ***Two-Way Interactions*** | | | | |
| Temperature × SM | 0.129 | 0.091 – 0.166 | <0.0001 | **** |
| Temperature × Sterol | −0.014 | −0.052 – 0.024 | 0.447 | ns |
| SM × Sterol | 0.033 | −0.005 – 0.071 | 0.088 | ns |
| ***Three-Way Interaction*** | | | | |
| Temperature × SM × Sterol | 0.004 | −0.050 – 0.058 | 0.873 | ns |

**Regression equation:**

$$GP = 0.433 - 0.154(Temp) - 0.027(SM) - 0.031(Sterol) + 0.129(Temp \times SM) - 0.014(Temp \times Sterol) + 0.033(SM \times Sterol) + 0.004(Temp \times SM \times Sterol)$$

Significance: **** $p < 0.0001$; * $p < 0.05$; ns, not significant. Dependent variable: GP ( laurdan generalized polarization). Least-squares multiple linear regression. SM, sphingomyelin; Erg, ergosterol; Temp, temperature.

**Table S1. Further details of ANOVA analysis of factors influencing membrane order.** Three factor ANOVA evaluating the effects of temperature, sphingolipid chain length (eSM vs. C26-SM), and sterol structure (cholesterol vs. ergosterol) along with their interactions on the measured response variable. P-values for each term are shown.

| Strain | Genotype/Description |
|---|---|
| W303a | MATa ura3-52 trp1Δ2 leu2-3_112 his3-11 ade2-1 can1-100 |
| *elo2*Δ | W303a, *elo2*Δ::*LEU2* |
| *elo3*Δ | W303a, *elo3*Δ::KanMX |
| CHOL | W303a, *erg6*Δ::*TRP1*-$P_{TDH3}$-DHCR7, *erg5*Δ::*LEU2*-$P_{TDH3}$-DHCR24 |

**Table S2.** List of yeast strains used in this study and their genotypes.
